# Comparative Metagenomics Provide Mechanistic Insights into the Biodegradation of the Non-hydrolysable Plastic Polyvinyl Chloride in Gut Microbiota of Insect Larvae

**DOI:** 10.1101/2024.02.06.579071

**Authors:** Haoran Peng, Zhe Zhang, Xiaoxi Kang, Yunhua Zhang, Huilin Zhang, Yuxuan Wang, Dongchen Yang, Jinlin Zhang, Yajie Wang, Yong-Guan Zhu, Feng Ju

## Abstract

Using microbiomes to mitigate global plastic pollution is of paramount importance. Insect microbiomes have garnered emerging interest for their ability to biodegrade non-hydrolysable plastic polymers. The larvae of *Spodoptera frugiperda*, a globally prevalent migratory crop pest, are accidentally discovered to consume polyvinyl chloride (PVC) plastic films, highlighting the role of their gut microbiome. Following the migration of *S. frugiperd* in China, this study displays a comprehensive geographical profile of its larval gut microbiota and a significant shift after PVC feeding. Using comparative metagenomics we revealed the functional redundancy within the larval gut microbiomes of two distinct insects after PVC ingestion, we discovered a surprisingly potent PVC-dechlorinating activity of an NADH peroxidase (6.48 mg/L chlorine produced in 96 hours with NADH as a cofactor) encoded by *Enterococcus casseliflavus* EMBL-3. These findings open a new avenue for understanding plastic biodegradation mechanism and enable the development of biotechnologies to mitigate global plastic pollution.

## Introduction

The ever increasing of synthetic plastic waste has emerged as an urgent and widespread global environmental challenge. Since 1950, the total global production of plastics has soared to a staggering 7.8 billion tons^1^. Regretfully, a substantial 79% of plastics have accumulated in the natural environment, with polyvinyl chloride (PVC) comprising 10% of the overall plastics usage ^2^. At present, conventional disposal methods for PVC waste predominately involved landfill and incineration. However, these methods generate secondary pollutants such as hydrogen chloride and dioxin, along with the emission of greenhouse gases ^3^. In light of these issues, alternative strategies like biological treatment and recycling or upcycling have emerged as effective and environmentally friendly pathways for addressing PVC plastic wastes and mitigating. These sustainable practices are crucial for a greener and more resilient future^4^.

Plastic polymers can be classified into hydrolysable plastics with C-O backbones (e.g., PET and PUR) and non-hydrolysable plastics with C-C backbones (PE, PP, PS and PVC)^5^. Remarkable progress has been made in the biodegradation of hydrolysable plastics, exemplified by the identification of the PET-degrading bacterium *Ideonella sakaiensis*^6^. Subsequent research has focused on the PET-degrading enzymes PETase and MHETase and the modification of the PET-degrading enzymes ^7,8^, eventually leading to significant strides towards large-scale PET biodegradation and recycling. In contrast, our understanding of the biodegradation of non-hydrolysable plastics remains limited. While various strains of fungi (e.g., *White Rot Fungi*^9^ and *Phanerochaete chrysosporium*^10^) and bacteria (e.g., *Pseudomonas citronellolis* and *Bacillus flexus*^11,12^) have been identified, most studies have primiarily reported the morphological and physicochemical changes of PVC plastic (e.g., weight loss and molecular weight). Notably, many studies have demonstated the ccapacity of insect gut microbiota to biodegrade plastics, including PS^13,14^, PET^15^, PVC^16,17^. However, little is known about the PVC-degrading genes and enzymes within these degrading strains or microbial communities^18^. Consequently, it is imperative to identify additional PVC-degrading enzymes and microbes and deepen our comprehesion on their biodegradation mechanisms.

To date, numerous studies have extensively documented the impacts of widespread plastic pollution on both the host gut microbiome and non-host associated environmental microbiome^19^. Certain insect species, particularly the larvae of darkling beetles, wax moths and meal moth, have garnered attention due to their remarkable ability to utilize and degrade various plastic polymers, such as polyvinyl chloride (PVC), polystyrene (PS) and polyethylene (PE) ^3,20,21^. This has led to widespread speculation regarding the symbiotic relationship between insects and their gut microbiomes, suggesting a crucial role in the adaptation of insects to environmental stress. In this process, a variety of microbes are believed to participate in the decomposition and elimination of xenobiotic pollutants such as biocides and plastics^22^. However, most research in this area has been confined to structural changes in microbial communities and identification of potential plastic degraders, lacking experimental validation. The connection between gut microbiota and plastic degradation mechanisms has been rarely explored.

Recently, we discovered that the larvae of the globally invasive agricultural pest, *Spodoptera frugiperda*, can subsist solely on PVC film. Employing multi-omics analysis, a gut bacterium *Klebsiella variicola* EMBL-1 was isolated and identified to participate in PVC depolymerization^18^. This study aims to unveil the PVC-degrading ability of the entire gut microbiota of *S. frugiperda* larvae. We first characterize the geographical influence on the gut microbiota of *S. frugiperda* larvae. Utilizing a comparative metagenomic approach, we then identify PVC-degrading genera and enzymes class, along with relevant biodegradation pathways. Furthermore, we successfully cultured and validated the PVC degradation capability of *Enterococcus casseliflavus* EMBL-3. This strain demonstrated remarkable dechlorination activity through NADH peroxidase, marking the first instance of such enzymatic activity in all polymers. These findings not only enhance the exploration of microbial and enzymatic resources available for PVC biodegradation, but also lay a foundation for in-depth investigation into the underlying mechanisms of this process.

## Results

### Characterizing microbiota in *S. frugiperda* larvae and surrounding soils from agricultural field

To unravel the gut microbiota within S. frugiperda larvae and their associated environmental niches, we systematically sampled and analyzed materials from several ecologically significant sources: bulk soils (BS) from cornfields impacted by the larvae, the larvae’s excreted feces (EF), and gut contents, encompassing intestinal feces (IF) and intestinal mucosa (IM), from active 3-5 instar larvae. Samples were obtained from cornfields spanning four provinces in China, ensuring a representative dataset.

We conducted extensive microbial community structure analyses of these collected samples, emphasizing both compositional and diversity metrics. Preliminary results illustrating the microbial community structures of the larval and soil samples are presented in Fig. 1 and Supplementary Fig. S1. Notably, the phylum-level microbial composition of larval gut samples (Supplementary Fig. S1) revealed a dominance of *Firmicutes* (IM: 66.9%±14.6%, IF: 50.6%±22.7%) and *Proteobacteria* (IM: 30.7%±15.1%, IF: 48.2%±22.5%). In contrast, the excreted feces (EF) primarily consisted of *Proteobacteria* (51.6%±23.3%) and *Bacteroidota* (46.8%±23.4%). Soil samples exhibited a more diverse microbial spectrum, including *Proteobacteria*, *Actinobacteriota*, and *Bacteroidota*, resulting in higher observed microbial richness and diversity indices compared to larval samples (Supplementary Fig. S2).

**Fig. 1.**
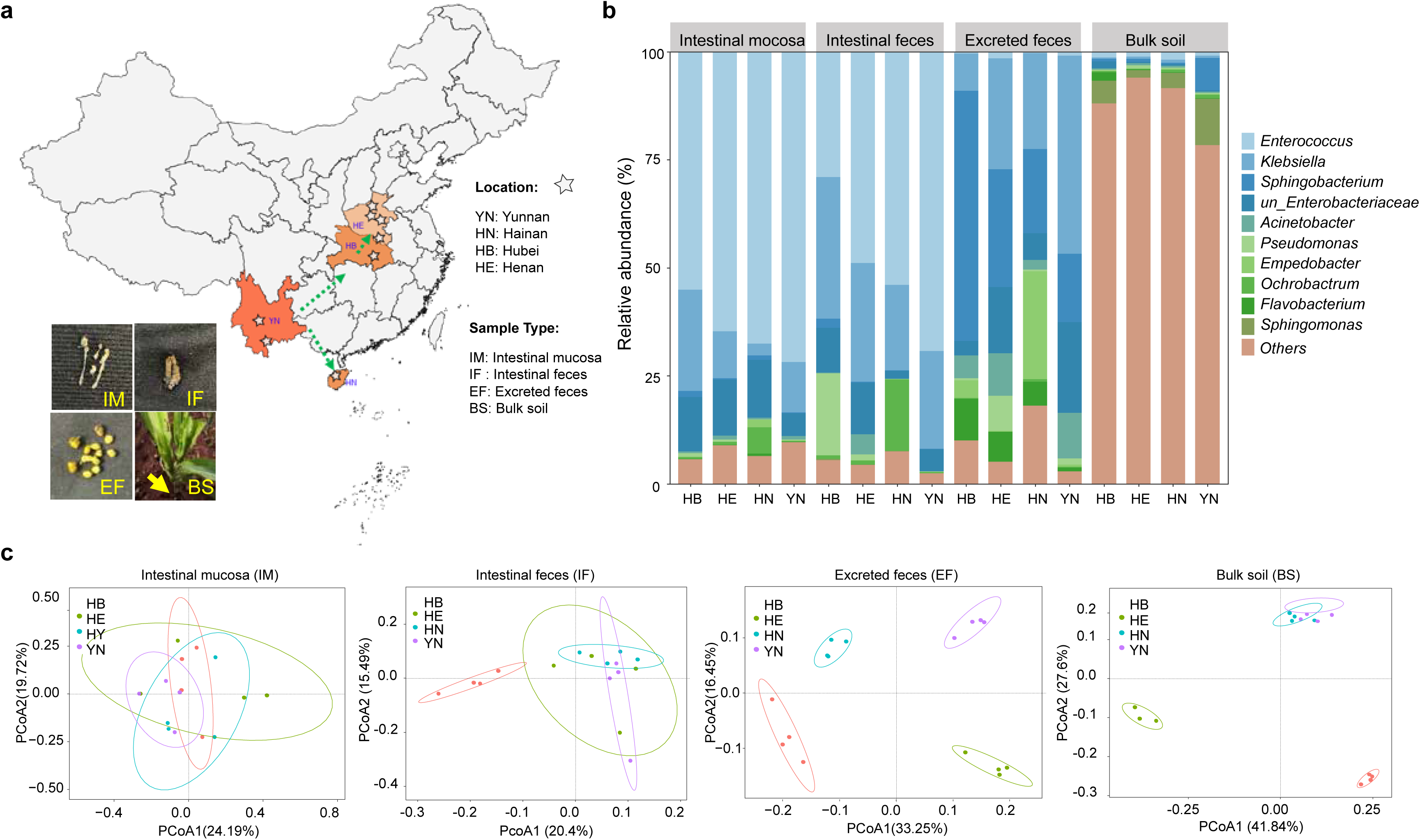
Field sampling and sequencing analysis results of collected samples of *S. frugiperda*. **a.** Location distribution map of field sampling of *S. frugiperda.* **b.** Species relative abundance maps of four types of samples collected in four provinces at the genus level. **c.** Beta-diversities of four types of samples collected in four provinces including Hubei (HB), Hebei (HE), Hainan (HN), Yunnan (YN). Unweighted-UniFrac PCoA was used to calculate the distance.

Moreover, microbial richness in larval gut samples was found to be highest in Hubei province, while the Hainan samples, located in a tropical monsoon zone, exhibited the greatest diversity (Supplementary Fig. S2). The larval gut was predominantly colonized by *Enterococcus* and *Klebsiella*, whereas *Sphingobacterium* and *Acinetobacter* were predominant in the excreted feces, indicating selective enrichment (Fig. 1b). Notably, microbial compositions of excreted feces and soil exhibited geographic differentiation (Fig. 1c). In contrast, the gut mucosa and feces microbiota showed a negligible geographical influence. This pattern suggests that the intestinal microbiota, primarily shaped by the host, remains consistent across regions, while the excreted feces and soil microbiota are influenced by local environmental factors. These findings provide an unprecedented glimpse into the microbial composition of *S. frugiperda* larvae, highlighting the influences of host biology and geographic factors. This study lays the groundwork for subsequent research into the gut microbiota of *S. frugiperda* larvae and its potential implications.

### PVC film degradation by larvae gut microbiota

In the experiment evaluating PVC degradation by *S. frugiperda* larvae, three groups were included: larvae fed with PVC film group (PVC group), larvae fed with Corn leaves group (Corn group), and larvae fed with PVC film and antibiotics group (Anti-PVC group). Previous studies measuring changes in physiological indexes supported the view that larvae can survive for days when fed with PVC film ^18^. To validate the effective depolymerization of PVC film, the film fragments in the PVC group and Anti-PVC group were collected to measure the changes in molecular weight using APC. The Mn of the PVC film fragments in the PVC group showed a significant (P-adj <0.05) increase by 24.6% compared with the control and Anti-PVC groups (Fig. S4a-S4b). This finding supported the notion that the degradation of PVC film by the *S. frugiperda* larvae is closely dependent on gut microbiota.

Moreover, GC-MS analysis demonstrated effective digestion of PVC film in the larval guts, resulting in the production of smaller organic molecules (Fig. S4c). In total, seven new substances (labelled as Compound 1 to 7) were detected in the PVC_F group but not the control group including 2-ethyl-1-hexanol, 2-ethyl-hexanoic acid, 2-nonanol, hexanedioic acid, 3-hydroxy-dodecanoic acid, hexadecenoic acid, and octadecanoic acid. Compounds 8 and 9 were identified as two kinds of known plasticizers, i.e., dioctyl adipate and dioctyl terephthalate. The PVC films were partially degraded during 5 days, encompassing the degradation of additives and PVC. Based on the structural formula of the compounds, compounds 1, 2, and 4 were likely derived from the degradation of plasticizers (compounds 8 or 9), and compounds 3, 5, 6, and 7 might be the potential PVC degradation products. This suggests that various components in PVC were degraded by multiple gut microbes through different pathways, indicating that the degradation of PVC films by the *S. frugiperda* larvae was from a combined action of diverse gut microbes.

### Comparative metagenomics of PVC-depolymerizing larvae gut microbiota of *S. frugiperda* and *T. molitor*

To unravel the microorganisms and enzymes contributing to PVC degradation and f facilitate the isolation, culturing, and functional analysis of PVC-degrading strains, we conducted comparative metagenomics between *S. frugiperda* and *T. molitor* larvae intestines.

#### Microbial diversity

In controlled indoor experiments, distinct microbiota responses to PVC ingestion. *S. frugiperda* larvae were observed. The larvae exhibited an increase in alpha-diversity of gut microbiota upon PVC consumption (Fig. 2a), contrasting with the decreased diversity observed in *T. molitor* larvae (Fig. 2c). Beta-diversity analyses revealed divergent microbial community structures between the PVC-fed and control groups in both species (Fig. 2b, Fig. 2d, S4d-S4e). Notably, PVC ingestion in *S. frugiperda* larvae led to a shift in gut and excreted microbial communities, with less dominant species becoming prevalent (Fig. S5a). Contrastingly, corn leaf feeding resulted in the dominance of *Enterobacteriaceae* (ASV1), suppressing other microbial species in *S. frugiperda* larvae, while PVC feeding disrupted this dominance, promoting the colonization of PVC-degrading bacteria such as ASV2 and ASV4 (*Enterococcus*) (Fig. S5a).

**Fig. 2.**
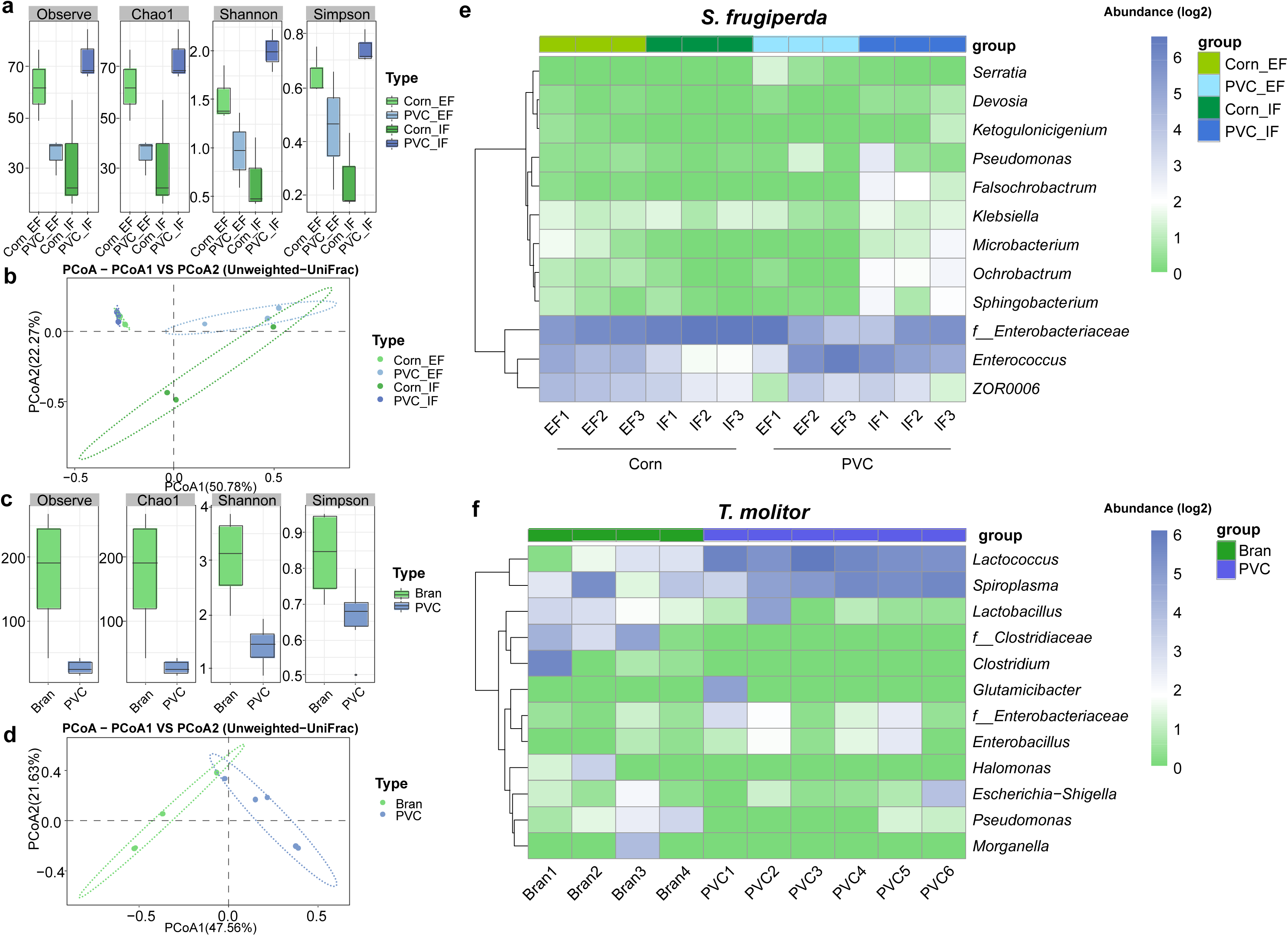
Microbial diversity and composition in *S. frugiperda* study and *T. molitor* study. **a.** Alpha-diversity, **b.** Beta-diversity, **c.** Alpha-diversity, **d.** Beta-diversity, **e.** Microbial composition in *S. frugiperda* study. **f.** Microbial composition in *T. molitor* study. Four alpha diversity indexes, observe, Chao1, Shannon, and Simpson were presented. And box plots showed center line as median, the upper and lower whiskers show maxima and minima, and box limits show upper and lower quartiles. Unweighted-UniFrac PCoA was used to calculate the distance, standardization method was Hellinger. The relative abundance of microbiota in each sample was presented in the heatmap, and log2 transfer was applied. The group annotation of samples was presented at the top of the heatmaps.

#### Microbial Composition

Further analysis revealed significant genus-level compositional shifts upon PVC feeding, with notable increases in genera such as *Enterococcus* (4.7±3.7% to 37.0±19.8%), *Ochrobactrum* (0.1±0.2% to 3.4±0.7%), *Klebsiella* (1.4±0.6% to 1.7±0.4%), *Falsochrobactrum* (0% to 2.80±1.40%), *Microbacterium* (0.03±0.05% to 2.23±1.55%) and *Sphingobacterium* (0.21±0.30% to 2.55±1.54%) (Fig. 2e), These shifts underscore a close link between gut microbiota composition and PVC biodegradation. Differential genus analysis identified distinctive microbial populations in PVC-fed larvae of both species, emphasizing the distinct microbial responses to PVC ingestion (Fig. 2f, Fig. S5b).

#### Metabolic Pathways

Despite the noticed compositional differences, the gut microbiota of *S. frugiperda* and *T. molitor* larvae shared functional similarities in PVC biodegradation. Metagenomic analysis revealed 593 genes significantly enriched in *S. frugiperda*’s PVC-fed group (Fig. 3a). Both species exhibited enrichment in eight key metabolic pathways related to biodegradation, such as other glycans degradation, phosphonates and phosphinate metabolism, and glycosaminoglycan degradation, highlighting functional convergence despite compositional variance (Figs. 3a-b).

**Fig. 3.**
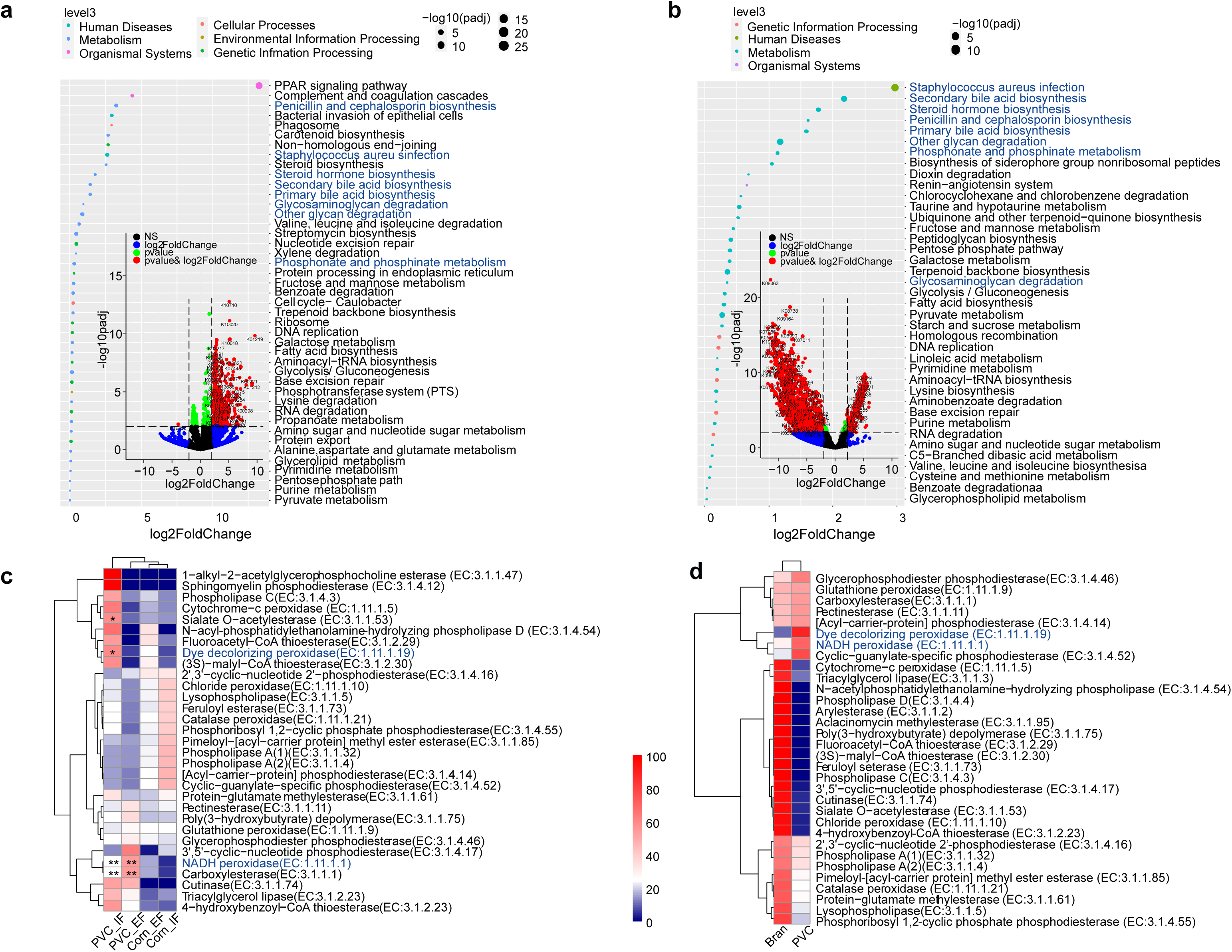
Differential enriched pathways and differential enriched enzymes. **a.** differential enriched genes and pathways in *S. frugiperda* study and **b.** in *T. molitor* study. (Volcano plot, cutoff: padj<0.01, log2FoldChange>2) **c.** Differential potential PVC degrading enzymes in *S. frugiperda* study **d.** and in *T. molitor* study. (pvalue<0.05, **pvalue<0.01). Based on differential abundance analysis of all genes, 57 potential plastic-degrading enzymes (currently reported in the literature and summarized and divided into five major categories, namely peroxidase, depolymerase, esterase, lipase, and cutinase) of intestinal microbiota were extracted, including 32 identified from *S. frugiperda* larvae and 33 from *T. molitor* larvae.

#### Degradation Enzymes

Plastic-degrading enzymes reported in the literature and their Enzyme Commission (EC) numbers were systematically summarized and divided into five major categories, namely peroxidase, depolymerase, esterase, lipase, and cutinase. Based on differential abundance analysis, 57 potential plastic-degrading enzymes were discovered, with significant enrichment of enzymes like sialate O-acetylesterase and carboxylesterase in PVC-fed groups (Fig. 3c). Interestingly, enzymes such as dye decoloring peroxidase and NADH peroxidase were enriched in PVC-fed groups across both species (Fig. 3d), suggesting their crucial role in PVC degradation. In *S. frugiperda*, dye decoloring peroxidase predominantly encoded in *Microbacterium* (ASV18), *Flavobacterium* (ASV33), and *Cellulosimicrobium* (ASV47), while NADH peroxidase was primarily encoded in *Enterococcus* (ASV2, ASV4) (Dataset S2). Conversely, in *T. molitor*, these enzymes were largely associated with ASVs belonging to *Glutamicibacter* and *Lactobacillus*, respectively (Dataset S3). These associations were particularly pronounced in one of the six parallel experimental groups. Despite potential issues with replication, this finding suggests the need to further investigate these bacterial genera for undiscovered PVC-degrading capabilities (Fig. 2f).

### In-depth characterization of population-level microbiota profiles with PVC biodegradation in *S. frugiperda* larvae

Building on our initial genus-level microbial composition findings, we delved deeper into the gut microbiota of *S. frugiperda* larvae, focusing on ASV-level abundance, particularly in relation to key enzyme carriers such as ASV2 and ASV4. This advanced analysis was driven by the need to understand the nuanced roles these specific microbial ASVs (as a proxy for population) play in PVC degradation.

Our investigation revealed a notable shift in the abundance of these key ASVs following PVC film ingestion. For instance, *Enterobacter*-affiliated ASV1, dominant in the gut microbiota under normal conditions, exhibited a marked decrease from 85.1±0.86% to 35.9±4.4%. In stark contrast, *Enterococcus*-affiliated ASVs (ASV2 and ASV4), identified as carriers of crucial plastic-degrading enzymes, displayed significant enrichment. ASV2’s abundance rose from 2.59±0.04% to 26.6±0.69%, while ASV4 increased from 1.91±0.01% to 10.3±0.13%. The more pronounced increase in ASV2 suggests its potentially superior role in PVC degradation.

Additionally, among the *Microbacterium* genus, all eight ASVs showed an increase in abundance, especially ASV12, which doubled from 0.2% to 0.4%. Similarly, within the *Sphingobacterium* genus, two branches displayed upward trends, most notably ASV11, which surged from 0.1% to 2.8% (Fig. 4). These findings not only reinforce our earlier observations but also establish a more detailed connection between specific ASVs, such as ASV2 and ASV4, and the potential for PVC degradation activity. This refined analysis at the ASV level provides a clearer picture of the intricate relationship between gut microbiota and plastic degradation in *S. frugiperda* larvae, enhancing our understanding of the microbial mechanisms involved in this process.

**Fig. 4.**
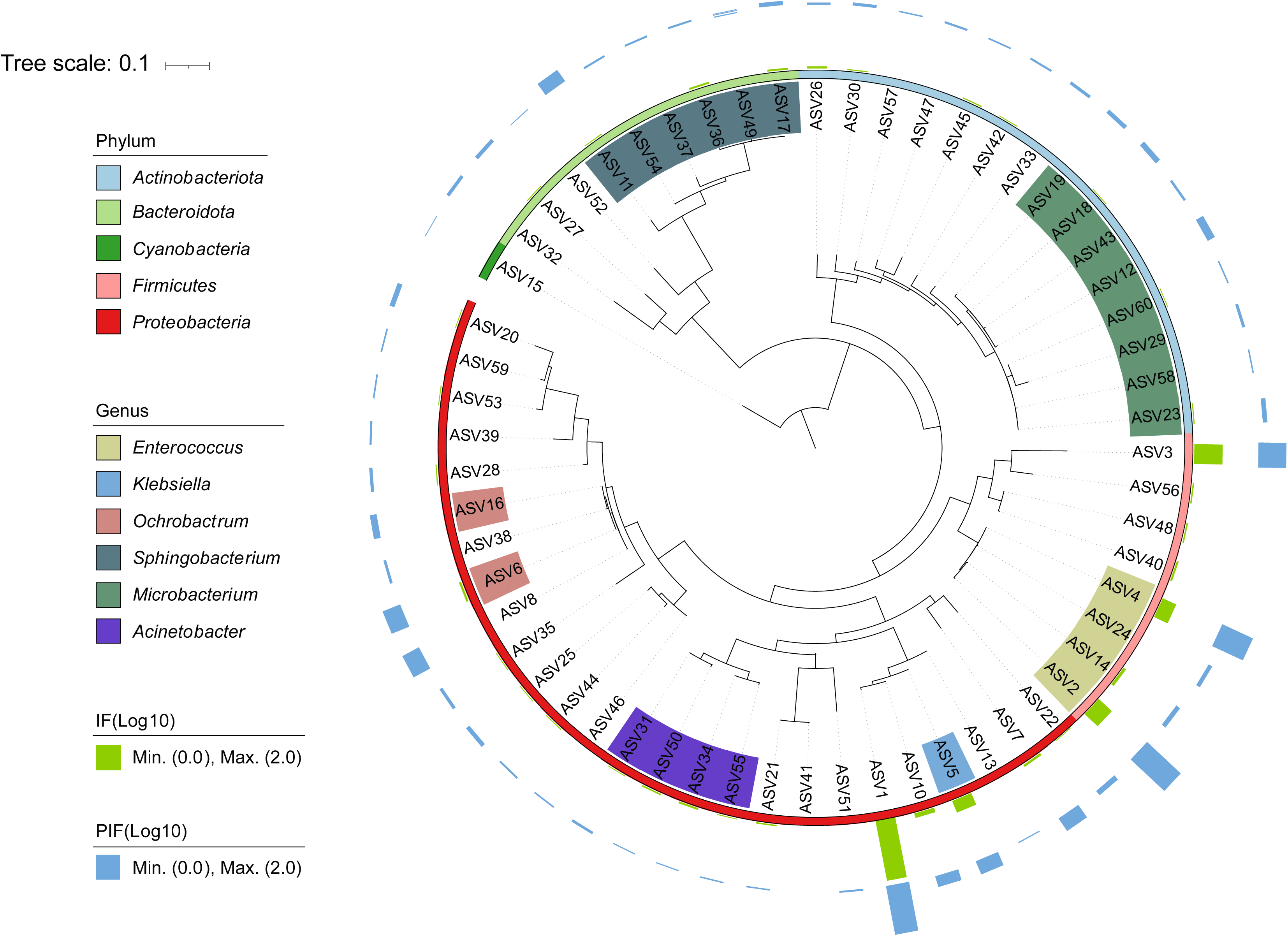
The phylogenetic tree and relative abundance annotation of the top 60 ASVs (Total relative abundance >97.5%) in *S. frugiperda*. Average relative abundances of the ASVs in corn-feeding groups were presented in green bars and PVC-feeding groups were presented in blue bars. Out layer presents the Phylum annotation and color labels on interested ASVs present the Genus level annotation.

### Targeted screening and functional verification of potential PVC-degrading strains

Following our comprehensive analysis, we identified microbiota enriched in PVC and possessing plastic-degrading enzymes. To isolate the potential PVC-degrading microorganisms and further validate NADH peroxidase function, the Known Media Database (KOMODO) was used to predict R2A as the appropriate medium for genus of *Enterococcus.* Through a series of screening and identification, one bacterial isolate was successfully separated from the genera *Enterococcus* (Fig. S6a). The NCBI’s online BLAST alignment of its full 16S rRNA gene sequence (Table S1) identified the isolate as *Enterococcus casseliflavus* (99.08%). Further comparison with ASVs showed *Enterococcus casseliflavus* was phylogenetically closer to ASV2, which significantly increased after PVC digestion (Fig. 5a). However, the experimental results of liquid culture using PVC film as the sole carbon source showed that the strain *Enterococcus casseliflavus* did not exhibited significant degradation activity (Fig. S6c).

**Fig. 5.**
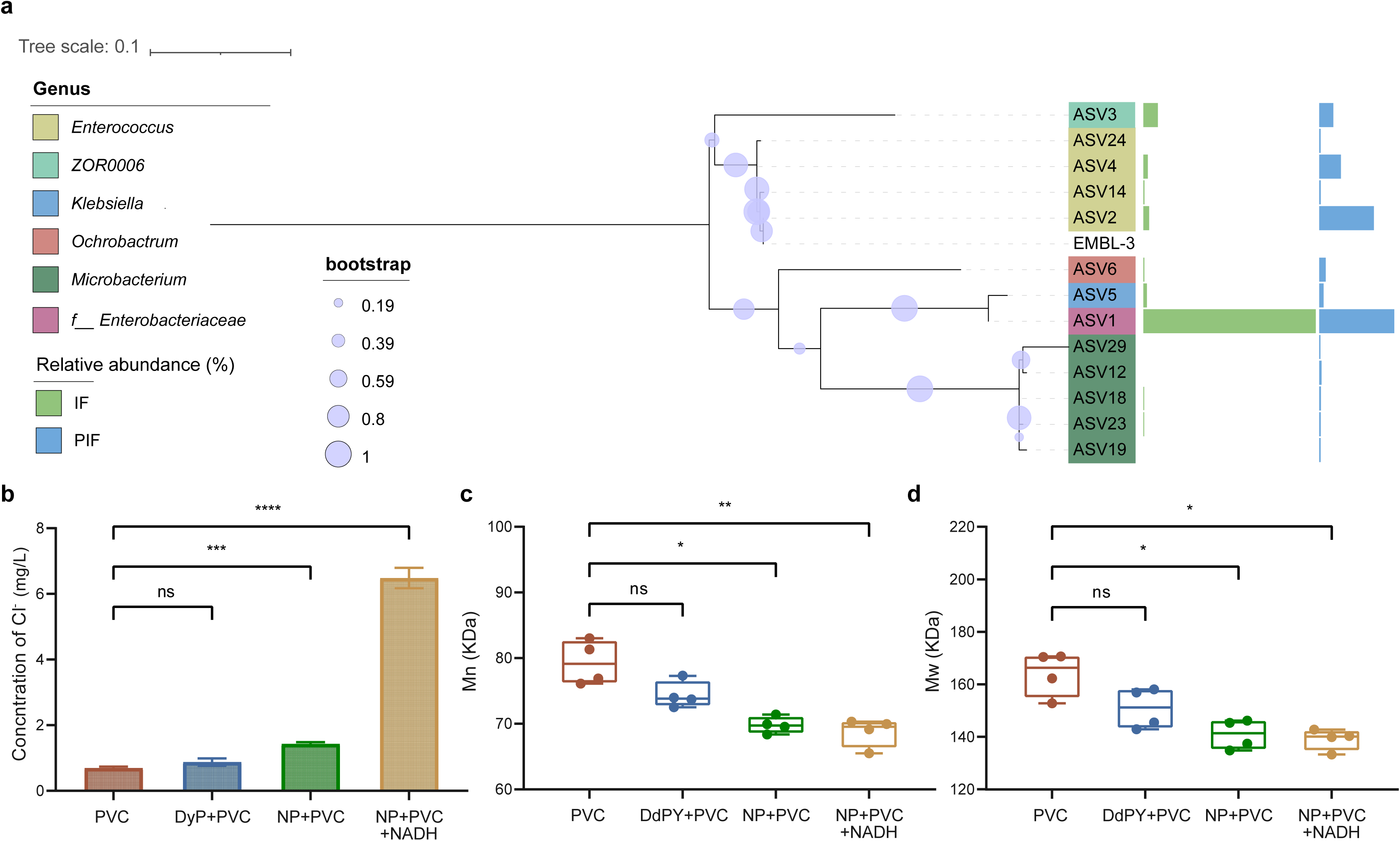
Functional verification of potential PVC-degrading enzymes. a, Phylogenetic tree of isolated strains and potential PVC degrading ASVs. Barplot on the right present the relative abundance of ASVs. b, Changes of chlorination concentration in the enzyme-treated supernatant (n = 3; Significance, p < 0.01 indicated by **, p < 0.001 indicated by ***, p < 0.0001 indicated by ****). c, Molecular weight (Mn) of PVC powder in the control group and the different enzyme treatment groups (n = 4; Significance, p < 0.01 indicated by **, p < 0.001 indicated by ***, p < 0.0001 indicated by ****). d, Molecular weight (Mw) of PVC powder in the control group and the different enzyme treatment groups (n = 4; Significance, p < 0.01 indicated by **, p < 0.001 indicated by ***, p < 0.0001 indicated by ****).

### Expression, purification, and functional verification of PVC-degrading enzymes

The comparative analysis between *S. frugiperda* and *T. molitor* in this study helped target two differentially degrading enzymes, i.e., dye decoloring peroxidase (EC:1.11.1.19) and NADH peroxidase (EC:1.11.1.1), and their bacterial host. The sequence of dye decoloring peroxidase could be found in our previous isolated strain EMBL-1(*Klebsiella* sp.)^18^, but the NADH peroxidase sequence could not. So, we obtained the pure strain of *E. casseliflavus* strain EMBL-3 and successfully extracted the NADH peroxidase sequence from its genome (Table S2). Then both enzymes were expressed and purified in vitro (Fig. S6b). Enzymatic assays confirmed that the dye decoloring peroxidase displayed peroxidase (1.23×10^5^ U/mg_prot_) activities and NADH peroxidase with peroxidase (9.8 U/mg_prot_) activities. Additionally, the dye decoloring peroxidase displayed dye decoloring activity (7.64%) to active blue 19.

To validate the degradation ability of these two enzymes on PVC, we set a 2 mL reaction system with 200 ug enzyme and 200 mg PVC powder, and measured molecular weight change and dechlorination of the PVC powder. The dye decoloring peroxidase displayed a feeble dechlorination on PVC powder (0.43 mg/L) (Fig. 5b). The APC result did not show significant change on the molecular weight (Mn decrease from 79.34 kDa to 74.37 kDa and Mw decrease from 164.04 kDa to 150.90 kDa) of PVC. In contrast, NADH peroxidase displayed a relatively stronger effect. As shown in Fig. 5c and 5d, the molecular weight of NADH peroxidase-treated PVC powder decreased significantly than those untreated. Mn decreased by 12.02% (from 79.34 kDa to 69.81 kDa) and Mw decreased by 14.07% (from 164.04 kDa to 140.97 kDa). This result indicated a depolymerization of PVC polymers chains by NADH peroxidase. A dechlorination of 0.74 mg/L was detected in the NADH peroxidase-treatment system compared to the control system (Fig. 5b). According to the putative structure and properties of the enzyme, the enzymatic activity of NADH peroxidase was presumed to increase in the presence of NADH. When 2 mM NADH was used as a substrate in combination with NADH peroxidase, a distinct dechlorination activity was detected, with the dechlorination amount detected increasing to 6.48 mg/L, while NADH alone did not have dechlorination activity on PVC (Fig. S6d). This indicates that NADH peroxidase may work on PVC polymer by dechlorination, making the first instance of dechlorination by enzymes in the pure PVC polymers. However, adding NADH did not have significant effect on molecular weight change of PVC. The inconsistencies in dechlorination changes and molecular weight changes made us aware that dechlorination contributes only partially to the polymer change. Therefore, we processed a series of mathematical calculations between the dechlorination amount and molecular weight change under several ideal assumptions (Method S9). We found that 6.48 mg/L dechlorination had almost no effect on molecular weight. In this case, it was speculated that dechlorination cannot explain the molecular weight change and there may be other depolymerizing reactions happen to the PVC polymer. Nevertheless, no degradation intermediates were detected in the reaction system.

## Discussion

Our research provides novel insights into the gut microbiota of an agricultural invasive pest *S. frugiperda* along its migratory pathways in China and reports, for the first time, a PVC biodegradation mechanism in the gut microbiota of *S. frugiperda* larvae. Our results suggest that the presence of functional redundancy, which is well known to induce stability of gut microbiota subjected to disturbance ^23^, in both *S. frugiperda* larvae and *T. molitor* larvae after PVC digestion. We identified a type of NADH peroxidase, possessed by two different microbes in two insect guts, which exhibited significant dechlorination ability. This enzyme was found to be encoded by the *Glutamicibacter* and *Lactobacillus* in *T. molitor* larvae gut, suggesting these two genera as potential PVC biodegradation resources. In *S. frugiperda* larvae gut, *E. casseliflavus* EMBL-3 encoded this enzyme.

The consistently identified differential enzymes, dye decoloring peroxidase and NADH peroxidase, were validated in the *Klebsiella* sp. Strain EMBL-1 ^18^ and *E. casseliflavus* strain EMBL-3. This marks the first discovery of dechlorination in PVC and polymers by enzymes. Microbial reductive dichlorination, a main route for biodegradation of chlorinated compound, involves replacing chlorine atoms with hydrogen atoms, requiring electron donors and energy^24,25^. Some bacteria in the environment, such as *Dehalococcoides* and *Dehalogenimonas* spp., were reported to have the capability of reductive vinyl chloride (VC) dechlorination to nontoxic ethene^25–29^. This study presumed the reductive dechlorination as the major biotransformation pathway of NADH peroxidase and dye decoloring peroxidase working on PVC, leading to increased free chloride, and decreased molecular weight. However, the degradation of PVC is a complicated process, in which the key steps, such as depolymerization, remains unrevealed. It may therefore be presumed that *Enterococcus sp.* act on additives in PVC film (such plasticizers), thus changing the physical and chemical properties of PVC film. These inferences require further verification through additional studies. Particularly, *E. casseliflavus*, intially thought to be closely associated with vegetation^30^, can be potentially acquired by vegetation-feeding pests. This species has been reported to degrade multiple substances including chlorantraniliprole, decabromodiphenyl ether, and a sulphonated azo-dye^31–33^. It well-documented capacity to dechlorinate compounds that are not polymers motivates us to further experimentally validate the PVC biodegradation ability of *E. casseliflavus* EMBL-3. However, we did not obtain direct experimental evidence of PVC biodegradation by EMBL-3, but instead successfully verified the enzymatic function of NADH peroxidase on PVC polymer. Considering the lack of observed PVC biodegradation by EMBL-3 in this study and the recently reported role of NADH peroxidase in preventing intracellular ROS formation in intestine bacteria ^34^, NADH peroxidase may be generated by the bacterium as a response to increased reactive oxygen species (ROS) induced by plastic stimuli, rather than being a dominant polymer degrader in EMBL-3.

Functional redundancy is prevalent in various animal gut microbiomes ^35,36^, including those of different species of insects. Our findings with plastic biodegradation in both *S. frugiperda* and *T. molitor* larvae well demonstrate that different insects can adapt and develop similar capacities to degrade pollutants, although they can have significantly different gut microbiota structure. While functional metagenomics was important for biodegradation function validation in those studies, shotgun metagenomics of gut microbiota was, however, hard to effectively perform in insect larvae due to its low microbial DNA content versus relatively high host DNA contamination. This outstanding technical bottleneck made 16S rRNA gene amplicon metagenomics being dominant method in insect larvae gut microbiota plastic degradation study^3,18,37^. Our study fully exploited the 16S rRNA gene amplicon metagenomics-based comparative analysis between insects and emphasized the importance of functional redundancy in the insect gut microbiota for plastic degradation. We believe differential analysis of microbial functions and enzymes can offer novel insights into functionally different microbes beyond abundance, as first demonstrated by our discovery of PVC polymer dechlorination by NADH peroxidase.

This research is the first to demonstrate the impact of geographical locations and host interactions on the gut microbiota of *S. frugiperda* larvae. Through a combination of dry and wet-lab analytical methods, we delved into microbial plastic degradation within the insect gut. Through comparative gut microbiota analysis, strain isolation, and culture enrichment techniques, validation of enzymatic mechanisms, we offer novel insights into potential PVC-degrading microbes, enzymes, and biodegradation mechanisms. These novel findings set the stage for future environmental research. Given the structural complexity of polymers like synthetic plastics, it is unlikely that a single enzyme or microorganism can facilitate complete degradation. Future studies should explore collaborative analyses of plastic degradation by various insect gut microbes and their enzymes, focusing on the synergistic activities of multiple microorganisms and enzymes in degradation processes.

## Materials and Methods

### Field sampling and processing of S*. frugiperda* specimen

According to the geographic spreading routes of *S. frugiperda* in China^38^, eight cornfields in eight cities of four provinces, i.e., Yunnan Province, Hainan Province, Hubei Province, and Henan Province, were selected for larvae and bulk soil sampling from during August 5^th^ and October 19^th^. Earlier studies on the gut microbes of the larvae of *Noctuidae* showed that the composition of gut microbes is relatively stable during the 3-5 instar larvae, such as the study of the gut microbial composition of the third and fifth instar larvae of *Spodoptera exigua* in the *Noctuidae* family through the sequencing of the V4 segment of 16S rRNA^39,40^. Therefore, the 3-5 instar larvae of *S. frugiperda* were selected for field and indoor experiments in this study. For each cornfield, 50 pieces of 3∼5 instar larval specimens of *S. frugiperda* were collected from the corn fields (Fig. 1a). For details on the experimental design plan for sample collection, see Method S1. The collected larvae were aseptically dissected in the evening, and the midgut of each larva was collected. Then the midgut was separated from the gut mucosa and gut contents by shaking and washing. The processing steps are described in Method S2. Samples name and locations could be found in Dataset S1. The excreted feces of the larvae were collected from 4 locations, including Yunnan (YN_EF), Hainan (HN_EF), Hubei (HB_EF) and Henan (HE_EF). Accordingly, the obtained four gut mucosa (IM) samples and four gut feces (IF) samples were respectively named: YN_IF, HN_IF, HB_IF and HE_IF; YN_IM, HN_IM, HB_IM and HE_IM, and the four samples of the bulk soil (BS) were named as YN_BS, HN_BS, HB_BS and HE_BS. All collected samples were stored in dry ice and sent back to the laboratory in 48 h, and then stored at -80 °C for further experiments. The undissected larvae specimens were stored at room temperature and fed with corn leaves.

### Indoor breeding of *S. frugiperda* and physicochemical characterization of PVC film

The larva of *S. frugiperda* were bred in an artificial climate room (temperature 22°C and humidity 55%) to explore the larval ability to live on PVC film, the specific experimental method can be found in our previous article^18^. The experimental design was briefly summarized as follows: 130 pieces of 4th-instar larvae with the same growth status were divided into four groups: 1) the control group (starvation, 15 pcs), 2) the Corn group (fed with corn leaves, 35 pcs), 3) the PVC group (fed with PVC film, 50 pcs), and 4) the Antibiotic group (fed with gentamicin sulfate antibiotic-soaked (100:3, *w*/*w*) corn leaves for 1 day before PVC film feeding).

The gut feces (IF) and excreted feces (EF) samples collected from the PVC and Corn groups were labelled PVC_IF, Corn_IF, PVC_EF and Corn_EF. Total DNA extraction and sequencing strategies were as previously described^18^. Fragments of PVC film in the excreted feces of the PVC groups (PVC_EF) were collected. Advanced Polymer Chromatography (APC, Waters, China) is used to detect the changes in the molecular weight of PVC films. For sample preparation and detection methods, see Method S3.

To obtain the products of PVC film biodegradation, PVC fraction in excreted feces (EF) in the Corn and PVC groups were collected for Gas Chromatography Mass Spectrometry (GC-MS) and named as Corn_F and PVC_F. Meanwhile, a control group (PVC film) was carried out and named as PVC_control. PVC fraction weighing 0.1 g was mixed with 10 mL of tetrahydrofuran, and the mixture was ultrasonicated for 30 min at room temperature. The extract was concentrated to 0.5 ml by drying with nitrogen gas and mixed with 1 mL of N-hexane to obtain some possible products by vortexing and ultrasonication for 10 min. The samples were filtered using a 0.22 µm PTFE syringe filter for subsequent steps^3,41^. The sample was injected at an initial temperature of 40 ℃ (hold, 4 min), which was progressively increased at 10 °C per minute and held at 280 °C (hold, 5 min). Moreover, the detector conditions, i.e., the transfer line temperature, ion source temperature, ionization mode electron impact and scan time, were maintained at 250°C, 280°C, 70 eV and 0.3 s, respectively.

### DNA extraction, PCR amplification and sequencing

Total DNA was extracted from gut feces samples collected using a QIAamp Fast DNA Stool Mini Kit following the manufacturer’s recommendations (QIAGEN GmbH, Germany). Then, the hypervariable V4-V5 regions of the prokaryotic 16S rRNA gene were amplified using 515F (5’-GTGYCAGCMGCCGCGGTAA-3’) and 926R (5’-CCGYCAATTYMTTTRAGTTT-3’). The modified 515F/926R double-ended primer was selected as the primer to amplify the V4/V5 segment. The amplicon products of each sample were evenly mixed and sequenced using a paired-end sequencing strategy (PE250) on the Illumina HiSeq2500 at Guangdong MAGIGene Technology Co., Ltd

### Bioinformatics, statistics and data visualization

#### Bioinformatics analysis

FastQC (Version 0.11.9) was used to check the quality of the obtained sequencing data. The double-ended data was imported through the software package Quantitative Insight into Microbial Ecology (QIIME2-2020.6), and DADA2 (1.14)^42^, which has higher sensitivity, more accurate clustering, and fewer false positives, was employed in construction of Amplicon Sequence Variation (ASVs). The qiime2 built-in package, the feature-classifier classify-sklearn machine learning method, was used for taxonomic classification using the SILVA SSU138^43^ as the reference database. To compare the larvae gut microbiome profile between this and prior studies, the SRA Toolkit (2.10.5) tool was used to download the 16S rDNA sequencing data of PVC degradation by *T. molitor* in the SRA database (download No. SRP186213)^3^. The experimental data of *T. molitor* includes PVC group (6 replicates) and Bran group (4 replicates). The data of indoor *S. frugiperda* larvae ingested the PVC film and the experimental data of *T. molitor* were processed by the same analysis method mentioned above.

### Statistical analysis and visualization

Processed data from Qiime2^42^ was imported into Rstudio Version 1.1.414 (R version 4.0.3). The phyloseq (1.32.0), MicrobiotaProcess, ggplot2 (3.3.3), and pheatmap (1.0.12) were used to do further analysis and visualization. R package Ggalluvial (v0.12.3) was used to conduct liquidity analysis^44^. Microbiota composition in Phylum-level were visualized by Sankey plots^44^. Lefse^45^ and Deseq2^46^ were used to perform differential analysis, and mafft^47^and fasttree^48^ were used to build the phylogenetic tree for the top 60 ASVs, default parameters were applied. The volcano plots were displayed with the R package Enhanced Volcano. PICRUST2^49^ was used to do microbial community function prediction. The abundances of genes and metabolic pathway were used for further analysis. Firstly, differential analysis was carried on genes, and differential enriched genes (padj<0.01, Log_2_Foldchange>2) were further mapped to pathways. Meanwhile, differential analysis was performed in all pathways, but only those differential genes mapped pathway were presented and considered to be the PVC enriched pathways in this study. All the above differential gene and pathway analysis was carried out in Deseq2^46^.

### Screening and functional verification of potential PVC-degrading strains and enzymes

The bioinformatics analysis results showed us that the bacteria of the genera *Sphingobacterium, Microbacterium, Enterococcus, Enterobacter, Ochrobactrum, Flavobacterium,* and *Cellulosimicrobium* were predicted to be related to PVC degradation. In order to further verify whether these bacteria of these genera have the activity of degrading PVC, we will adopt the method of isolation and screening to obtain and verify them. It is difficult to isolate and screen target strains directly from midgut microorganisms, so the Known Media Database (KOMODO)^50^ was used to select appropriate screening medium type for potential PVC-degrading strains. Based on the results of KOMODO predictions, a purposeful screening of bacteria was conducted. First, the gut feces obtained from the PVC group of *S. frugiperda* were diluted step by step (10^-1^ to 10^-8^) by the serial dilution method, and three solutions of 10^-5^ to 10^-7^ concentrations were selected for plating culture. All treatments were placed in a 30°C incubator for static culture. According to the basic characteristics of the target microorganism, some single colonies were selected from the plate for continuous culture until a single colony was obtained.

Based on the appearance and morphology of the strains, 16S rRNA method was used to amplify the full-length sequence for strain identification. The full-length sequence of the 16S rRNA gene of the strain was amplified with 27F and 1492R universal primers^51^, and then the sequences were aligned against the NCBI’s 16S ribosomal RNA sequences (Bacteria and Archaea) database using BlastN tool.

The obtained target strains were tested for the degradation activity of the self-made PVC film (Method S4). The verification system is an MSM liquid medium system with self-prepared pure PVC membrane as the only carbon source and energy source. All treatments were cultured at 30°C and 150 rpm for 42d, and the MSM medium was changed at 21d to maintain nutrient requirements for strain growth. Finally, the degradation activity of the strains was evaluated through the change of the molecular weight of the PVC film on 42d (Method S5).

### Expression, purification and functional verification of potential PVC-degrading enzymes

The bioinformatics analysis results showed us information about potential PVC-degrading enzymes. Two enzymes (dye decoloring peroxidase and NADH peroxidase) were analyzed to be potential PVC-degrading enzymes. Based on the genomic information of EMBL-1(*Klebsiella* sp.) and EMBL-3 (*Enterococcus* sp.), We extracted separately the gene information of dye decoloring peroxidase and NADH peroxidase from the genome of EMBL-1 and *Enterococcus* sp. strain and then expressed and purified these two enzymes in vitro by prokaryotic expression and purification method. The method of expression and purification of protein in vitro (Method S6) was used to obtain the pure enzyme for further enzyme activity assay (Method S7). Moreover, the method same as catalase-peroxidase^18^ was used to validate the PVC-degrading activity of the two enzymes (Method S7).

## Acknowledgements

This work was supported by the National Science Foundation of China (Grant No. 22241603 to JF), the Westlake Center for Synthetic Biology and Integrated Bioengineering (Grant No. WU2022A009 to JF), the Research Center for Industries of the Future (Grant No. WU2022A009 to JF), and the National Natural Science Foundation of China (grant no.: 32100091 to ZZ). JF also acknowledged support from the Biomedicine and the Research Center for Industries of the Future (WU2022C030) at Westlake University and the HRHI program 202309010 of Westlake Laboratory of Life Sciences. We thank Dr. Yinjuan Chen and Ke Wang from the Instrumentation and Service Center for Molecular Sciences at Westlake University for their assistance and discussion during the experiment. We also thank Dr. Xiaoyuan Zhao, Dr. Guoping Li, Dr. Shiyou Yan and Haixia Tian in different provinces for support and assistance during the field sampling.

## Data availability

The raw sequence data produced in this study has been uploaded to the NCBI under the accession numbers PRJNA877394, CNP0004265 and CNA0069186.

## Author contribution

F.J. obtained funding and supervised this study. F.J., H.P., Z.Z. and X.K conceptualized this study. Z.Z., H.P., D.Y., and J.Z. contributed to the collection of field larvae samples. H. P. performed metagenomics data analysis and strain analysis. Z.Z. and H.Z. performed strain culture. X.K and Y.X.W performed enzyme purification and degradation validation. Y.J. W. provided design and improvement suggestions to the enzyme verification experiment. H.P., Z.Z, X.K. and F.J. drafted the manuscript. Y.G.Z reviewed and edited the manuscript. F.J. edited and finalized the manuscript. All authors read and approved the final version of the manuscript.

---

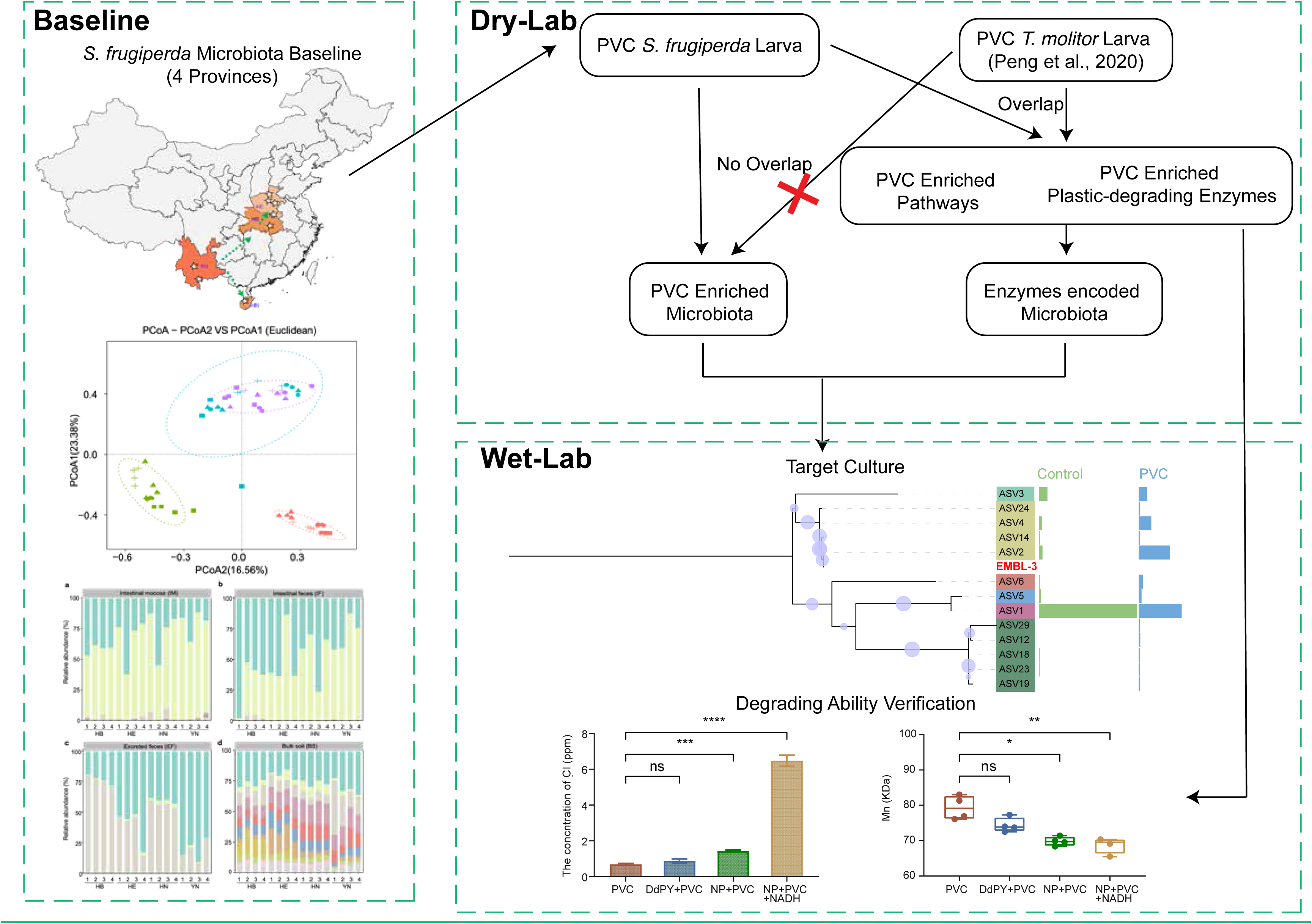

