## Supplemental Methods, tables and figures for "Comparative Metagenomics Provide Mechanistic Insights into the Biodegradation of the Non-hydrolysable Plastic Polyvinyl Chloride in Gut Microbiota of Insect Larvae"

### Supplementary Methods

**Method S1 Design plan for collecting samples of *S. frugiperda* in the field**

**Method S2 Methods of dissection of larvae and methods of separating intestinal feces and intestinal mucosa**

**Method S3 Preparation and detection methods of Advanced Polymer Chromatography**

**Method S4** **Preparation method of self-made PVC film**

**Method S5 The detection methods of molecular weight of the PVC film**

**Method S6 The method of expression and purification of protein in vitro**

**Method S7 The functional verification of potential PVC-degrading enzymes**

**Method S8 Detection method of chloride**

**Method S9 Mathematical calculations between the dechlorination and molecular weight**

**Table S1 The full-length 16S rRNA sequence of the strains screened from the intestinal microbes**

**Table S2 The gene information of dye decoloring peroxidase from strain EMBL-1 and NADH peroxidase from strain EMBL-3**

**Fig. S1 Phylum-level relative abundance of samples from four provinces**

**Fig. S2 Alpha-diversity of samples from four provinces**

**Fig. S3 Beta diversity of samples from four provinces collected in four sites**

**Fig. S4 The Degradation characterization of PVC and composition of intestinal microbes after larvae ingested PVC.**

**Figure S5 Overlapped microbiota and differential microbiota between samples in S. frugiperda study**

**Fig. S6 Functional verification results of potential PVC-degrading strains and enzymes**

### Supplementary Methods

### Method S1 Design plan for collecting samples of *Spodoptera frugiperda* in the field

**Bulk soil** Four plots were collected in each province, five sites were collected from each plot, and 100g of corn plant bulk soil was collected from each site. Use a sterile shovel to collect the bulk soil and store it in a sterile ziplock bag. The samples from 5 sites are mixed and marked for next step. The soil samples were stored in a 4°C refrigerator, and then stored in -20°C for use.

***S. frugiperda* lavae** Four plots were collected in each province, and 50 pieces of 3-5 instar active larva were collected from each plot, and a small amount of corn leaves were picked at the same time for normal feeding. The collected larval specimen was then placed in a ventilated carton for temporary storage until being dissected in the laboratory in 12 h, and then the guts were stored at -80°C until further operation and DNA extraction.

**Excreted feces** Since the feces of the larvae in the field were scattered, we collected the feces of the larva and placed them in the tubes. The collected feces samples were stored at 4°C for temporary storage until they were transported to the laboratory in 12 h, and then they were stored at -80°C until further operation and DNA extraction.

### Method S2 Methods of dissection of larvae and methods of separating intestinal feces and intestinal mucosa

Place the larvae on ice to weaken their activity, put the larvae in a 2ml centrifuge tube containing 70% alcohol for 2 minutes, and then transfer them to sterile water rinse 3 times(each time for 1 minute). Then clamp the head of the larva, make a small mouth under the head, cut down from the small mouth along the body of the larva, gently grip the larvae’s intestines with tweezers and pull them out, and place them in a sterile 1.5ml centrifuge tube for later use. Cut the intestine into two sections with scissors, shake and wash the intestine 3 times with PBS solution, pick out the intestinal mucosa, and merge the washing liquid to form the intestinal suspension (Intestinal Feces). Merge the selected intestinal mucosa, shake and wash them 3 times with PBS solution, and merge the washing solution to obtain the intestinal mucosal solution (Intestinal Mucosa).

### Method S3 Preparation and detection methods of Advanced Polymer Chromatography

PVC fraction (0.2 g) from the feace were first extracted with Tetrahydrofuran (20 mL). After filtration, the solution was mixed on a magnetic stirrer with gentle heating (60 ◦C). After the extracted solution was concentrated to a total volume of 5 mL^1^, the extract (60 μL in volume) was injected into the Advanced Polymer Chromatography (APC) measurements.

Advanced Polymer Chromatography (APC) measurements were carried out with three columns XT900-XT450-XT 200 (2.5um, 4.6*150mm) and a RI detector. Tetrahydrofuran was used as mobile phase (0.4 ml/min) after calibration with polystyrene standards of known molecular mass. Untreated PVC film was used as control group.

### Method S4 Preparation method of self-made PVC powder and PVC films

Because the PVC film purchased (Sinopec Yanshan petrochemical company) contains too many additives, we chose to prepare pure PVC powder and film using the purchased PVC film by extraction method. The preparation method of the pure PVC film referred to by the national standard method (GB/T 39110-2020) with appropriate modifications, as follows: PVC film (1 g) were dissolved in 30 ml of tetrahydrofuran. 70 ml of methanol solution was added to the solution and left to stand until PVC was precipitated. The PVC pellet was obtained by centrifugation and rinsed three times with methanol. The cleaned PVC pellet were places in a fume hood at room temperature for 2 days to thoroughly evaporate the methanol. At last, the cleaned PVC precipitate was collected and dried at 45°C for 24h, which was pure PVC powder. Then the PVC powder was dissolved in 30 ml of tetrahydrofuran again and then this solution (6 ml) was dropped on a glass Petri dish (Φ 90 mm). The solvent was allowed to evaporate and a film was formed. The PVC films were placed in a fume hood at room temperature for 2 days to thoroughly evaporate the dichloromethane. The PVC films were sterilized with 75% ethanol (v/v) and rinsed with sterile water. They were then cut into sheets of 30 × 30 mm square for the growth of bacterial suspension in a liquid medium respectively.

### Method S5 The detection methods of molecular weight of the PVC film

Advanced polymer chromatography (APC, Waters, China) measurements were carried out with three columns XT900-XT450-XT 200 (2.5um, 4.6*150mm) and a RI detector. Tetrahydrofuran (THF, HPLC) was used as mobile phase (0.4 ml/min) after calibration with polystyrene standards of known molecular mass. Non-incubated PVC film was used as references.

### Method S6 Method of expression and purification of protein in vitro

Based on previous research results in the laboratory. We extracted the gene information of dye decoloring peroxidase and NADH peroxidase from the genome of EMBL-1 and EMBL-3 strain, expressed and purified these two enzymes in vitro by prokaryotic expression and purification method.

Appropriate restriction sites were added to the primers (F: A**GGATCC**ATGTCTCAGGTTCAGAGCG, R: A**CTCGAG**CAGCGCCTGAATACGCTCCA, the bold font showed the restriction endonuclease sites BamHI and XhoI) designed to correspond to dye decoloring peroxidase. While the primers (F: GG**GGATCC**ATGAAAATCATTATCATAGGCGGG, R: AT**CTCGAG**AGTCTCATCGGCATCACTTC, the bold font showed the restriction endonuclease sites BamHI and XhoI) to NADH peroxidase.

Primer synthesis and DNA sequencing services were provided by Shanghai Bioengineering Co., Ltd.(Shanghai, China). Using procedures developed earlier^2^, we constructed the expression vector and conducted the expression and purification in vitro. E.coli (DE3) cells containing the constructed vectors were inoculated into fresh LB medium containing kanamycin (0.5 mg/mL) and incubated at 37°C in a rotary shaker at 150 rpm until reaching an OD600 of 0.6. The recombinant strains were then induced with 0.6 mM isopropyl-b-D-thiogalactopyranoside (IPTG) at 37 °C for 2 h. The cells were collected by centrifugation at 6000 rpm for 5 min, and target protein expression was verified by 10% SDS-PAGE. The cells were resuspended in buffer A (20 mM Tris-HCl, 300 mM NaCl, 0.1% Triton-100, pH 8.0) in an ice bath, lysed by ultrasonication and centrifugation at 12000 rpm for 20 min. The supernatant was purified using a Ni-IDA agarose magnetic beads (Beijing Biomed Co., Ltd.) as the manufacturer's instructions. The eluate was dialyzed against 20 mM Tris, 50 mM NaCl, pH 8.0, and the purity of the recombinant proteins was determined by 10% SDS-PAGE. Protein concentrations were determined by the Bradford method.

### Method S7 Functional verification of potential PVC-degrading enzymes

We determined the activities of peroxidase (POD) of dye decoloring peroxidase and NADH peroxidase by using kits purchased from Beijing Solarbio Science (China)，and all measurements were performed according to the manufacturer’s instructions. Moreover, the dye decolorization activity of peroxidase was also tested with active blue 19 (RB19) according to the former method^3^.

Peroxidase (POD) activity determination: POD catalyzes H_2_O_2_ to oxidize specific substrates and has characteristic light absorption at 470nm. Definition of unit: 0.01 change in A470 per minute per mg of protein per ml of reaction system is an enzyme activity unit.

Dye decolorizing assay: RB19 (final concentration ) was dissolved in 50 mM sodium citrate buffer (pH 4.5). Dye decolorization peroxidase (final concentration 1 mg/mL) and H_2_O_2_ (final concentration 1 mM) were added to initiate the reaction. The decolorization of RB19 was monitored at 595 nm at 0, 5, 10, and 30 min. The decolorizing activity (DA) was calculated based on the following equation.

$$\text{DA(\%)=}\frac{A_{i}-A_{f}}{A_{i}}\times100$$

where *A_i_* and *A_f_* are the initial and the final absorbance, before and after treatment.

We also designed an assay to detect the degradation activity of the dye decoloring peroxidase and NADH peroxidase on pure PVC. The experimental design was as follows: 1) Reaction Buffer (20 mM phosphate buffer, 2mL) + dye decoloring peroxidase (100 ug/mL), 2) Reaction Buffer (20 mM phosphate buffer, 2mL) + dye decoloring peroxidase (100 ug/mL)+PVC (200 mg), 3) Reaction Buffer (20 mM phosphate buffer, 2mL) +PVC (200 mg)，4) Reaction Buffer (20 mM phosphate buffer , 2mL) + NADH peroxidase (100 ug/mL), 5) Reaction Buffer (20 mM phosphate buffer, 2mL) + NADH peroxidase (100 ug/mL)+PVC (200 mg), 5) Reaction Buffer (20 mM phosphate buffer, 2 mL) + NADH peroxidase (100 ug/mL) + PVC (200 mg) + NADH (0.02, 0.2, 2 mM), 6) Reaction Buffer (20 mM phosphate buffer, 2 mL) + PVC (200 mg) + NADH (0.02, 0.2, 2 mM), 7) Reaction Buffer (20 mM phosphate buffer, 2 mL) + NADH peroxidase (100 ug/mL) + NADH (0.02, 0.2, 2 mM). All treatments were repeated three times and cultured in a shake (150 rpm, 30℃) for 96 h.

### Method S8 Detection method of chloride

The liquid sample was filtered through a 0.22-μm membrane filter before measurement. A Dionex Aquion RFIC ion chromatograph system (Thermo Fisher Scientific, Inc.) was used. The mobile phase was KOH, the flow rate was set to 0.30 ml/min and the separation was performed on a Dionex Ion Pac AS18-Fast analysis column (2*150 mm; Thermo Fisher Scientific, Inc.), and attached a Dionex Ion Pac AG15 anion guard column (2*250 mm; Thermo Fisher Scientific, Inc.). The column temperature was maintained at 30˚C during the operation. The injection volume was 25 µl. In this experiment, 5, 40 and 5 mM mobile phase (KOH) were used.

### Method S9 Mathematical calculations between the dechlorination and molecular weight

Assume that all polymer molecules have the same molecular weight.

For number-average molecular weight:

Initial number-average molecular weight

$$M_{n0}= 79.34 kDa = 7.9340\times{10}^{4} g/mol$$

Total number of PVC molecule

$$n_{tn}=\frac{m_{PVC0}}{M_{n0}}\times NA=2.5208\times{10}^{-6}NA$$

PVC weight after reaction

$$m_{PVC}'=m_{PVC0}-c_{deCl}\times V=0.1999 g$$

Number-average molecular weight after reaction

$$M_{n}^{'}=\frac{m_{PVC}'}{n_{tn}}=7.9335\times{10}^{4} g/mol=79.335 \mathrm{kDa}$$

For mass-average molecular weight:

Initial mass-average molecular weight

$$M_{w0}= 164.04 kDa = 1.6404\times{10}^{5} g/mol$$

Total number of PVC molecule

$$n_{tw}=\frac{m_{PVC0}}{M_{n0}}\times NA=1.2192\times{10}^{-6}NA$$

Number of brackets in one PVC molecule

$$b=\frac{M_{w0}}{62.5 g/mol}=2.6246\times{10}^{3}$$

Number of chloride

$$n_{Cl}=\frac{c_{deCl}\times V}{M_{Cl}}\times NA=3.6507\times{10}^{-7}NA$$

Assume all chloride were removed in one molecule.

Molecular weight after chloride were removed

$$M_{w/o-Cl}=M_{w0}-b\times\left( M_{Cl}-M_{H} \right)=7.3490\times{10}^{4} g/mol$$

Number of molecules without chloride

$$n_{w/o-Cl}=\frac{n_{Cl}}{b}=1.3909\times{10}^{-10}NA$$

Number of molecules without dechlorination

$$n_{w/-Cl}=n_{tw}-n_{w/o-Cl}=1.2191\times{10}^{-6}NA$$

Mass-average molecular weight after reaction

$$M_{w}^{'}=\frac{M_{w/o-Cl}\times n_{w/o-Cl}+M_{w0}\times n_{w/-Cl}}{n_{tw}}=1.6403\times{10}^{5} g/mol=164.03 \mathrm{kDa}$$

Assume only one chloride was removed in one molecule.

Molecular weight after chloride were removed

$$M_{deCl}=M_{w0}-\left( M_{Cl}-M_{H} \right)=1.6401\times{10}^{5} g/mol$$

Number of molecules that one chloride was removed

$$n_{deCl}=n_{Cl}=3.6507\times{10}^{-7}NA$$

Number of molecules that does not change

$$n_{non}=n_{tw}-n_{deCl}=8.5414\times{10}^{-7}NA$$

Mass-average molecular weight after reaction

$$M_{w}^{''}=\frac{M_{deCl}\times n_{deCl}+M_{w0}\times n_{non}}{n_{tw}}=1.6403\times{10}^{5} g/mol=164.03 \mathrm{kDa}$$

1	Peng, B. Y. *et al.* Biodegradation of Polyvinyl Chloride (PVC) in Tenebrio molitor (Coleoptera: Tenebrionidae) larvae. *Environ Int* **145**, 106106, doi:10.1016/j.envint.2020.106106 (2020).

2	Zhang, Z., Zhang, Y., Yang, D. C. & Zhang, J. L. Expression and functional analysis of three nicosulfuron-degrading enzymes from Bacillus subtilis YB1. *J Environ Sci Health B* **53**, 476-485, doi:10.1080/03601234.2018.1455344 (2018).

3	Dhankhar, P. *et al.* Characterization of dye-decolorizing peroxidase from Bacillus subtilis. *Arch Biochem Biophys* **693**, 108590, doi:10.1016/j.abb.2020.108590 (2020).

### **Table S1** the full-length 16S rRNA sequence of the strain screened from the gut

| **Name** | **full-length 16S rRNA sequence** |
| --- | --- |
| EMBL-3  (*Enterococcus)* | AGAGTTTGATCCTGGCTCAGGACGAACGCTGGCGGCGTGCCTAATACATGCAAGTCGAACGCTTTTTCTTTCACCGGAGCTTGCTCCACCGAAAGAAAAAGAGTGGCGAACGGGTGAGTAACACGTGGGTAACCTGCCCATCAGAAGGGGATAACACTTGGAAACAGGTGCTAATACCGTATAACACTATTTTCCGCATGGAAGAAAGTTGAAAGGCGCTTTTGCGTCACTGATGGATGGACCCGCGGTGCATTAGCTAGTTGGTGAGGTAACGGCTCACCAAGGCAACGATGCATAGCCGACCTGAGAGGGTGATCGGCCACACTGGGACTGAGACACGGCCCAGACTCCTACGGGAGGCAGCAGTAGGGAATCTTCGGCAATGGACGAAAGTCTGACCGAGCAACGCCGCGTGAGTGAAGAAGGTTTTCGGATCGTAAAACTCTGTTGTTAGAGAAGAACAAGGATGAGAGTAAAATGTTCATCCCTTGACGGTATCTAACCAGAAAGCCACGGCTAACTACGTGCCAGCAGCCGCGGTAATACGTAGGTGGCAAGCGTTGTCCGGATTTATTGGGCGTAAAGCGAGCGCAGGCGGTTTCTTAAGTCTGATGTGAAAGCCCCCGGCTCAACCGGGGAGGGTCATTGGAAACTGGGAGACTTGAGTGCAGAAGAGGAGAGTGGAATTCCATGTGTAGCGGTGAAATGCGTAGATATATGGAGGAACACCAGTGGCGAAGGCGGCTCTCTGGTCTGTAACTGACGCTGAGGCTCGAAAGCGTGGGGAGCGAACAGGATTAGATACCCTGGTAGTCCACGCCGTAAACGATGAGTGCTAAGTGTTGGAGGGTTTCCGCCCTTCAGTGCTGCAGCAAACGCATTAAGCACTCCGCCTGGGGAGTACGACCGCAAGGTTGAAACTCAAAGGAATTGACGGGGGCCCGCACAAGCGGTGGAGCATGTGGTTTAATTCGAAGCAACGCGAAGAACCTTACCAGGTCTTGACATCCTTTGACCACTCTAGAGATAGAGCTTCCCCTTCGGGGGCAAAGTGACAGGTGGTGCATGGTTGTCGTCAGCTCGTGTCGTGAGATGTTGGGTTAAGTCCCGCAACGAGCGCAACCCTTATTGTTAGTTGCCATCATTTAGTTGGGCACTCTAGCGAGACTGCCGGTGACAAACCGGAGGAAGGTGGGGATGACGTCAAATCATCATGCCCCTTATGACCTGGGCTACACACGTGCTACAATGGGAAGTACAACGAGTTGCGAAGTCGCGAGGCTAAGCTAATCTCTTAAAGCTTCTCTCAGTTCGGATTGTAGGCTGCAACTCGCCTACATGAAGCCGGAATCGCTAGTAATCGCGGATCAGCACGCCGCGGTGAATACGTTCCCGGGCCTTGTACACACCGCCCGTCACACCACGAGAGTTTGTAACACCCGAAGTCGGTGAGGTAACCTTTTGGAGCCAGCCGCCTAAGGTGGGATAGATGATTGGGGTGAAGTCGTAACAAGGTAACC |

### **Table S2** the the gene information of dye decoloring peroxidase from strain EMBL-1 and NADH peroxidase from strain EMBL-3

| **Name** | **gene sequence** |
| --- | --- |
| dye decoloring peroxidase | ATGTCTCAGGTTCAGAGCGGCATTTTGCCGGAACATTGCCGCGCGGCGATTTGGATTGAAGCCAATGTCAAAGGGGACGTTAACGCCCTGCGCGAAGCGAGCAAAAATTTTGTCGATAACGTGGCCACCTTCCAGGCTAAATTCCCCGACGCTAAACTCGGGGCGGTGGTGGCGTTCGGCAATAACGTCTGGCGTCAGCTGAGCGGCGGCGAAGGGGCGGAAGAGTTAAAAGATTTTCCGGTCTATGGCAAAGGGCTGGCGCCGTCCACCCAGTATGACCTGCTGATTCATATTTTATCCGCCCGCCATGAAGTTAACTTCTCGGTGGCGCAGGCCGCGATGGCTGCCTTTGGCGATGCTATCGACGTGAAAGAAGAGATCCACGGTTTCCGTTGGGTGGAAGAGCGTGATCTCAGCGGCTTCGTCGACGGCACCGAAAACCCGGCGGGGGAAGAAACCCGCCGCGAAGTGGCGGTCATTAAAGACGGTGTTGACGCGGGCGGCAGCTACGTGTTCGTCCAGCGCTGGGAGCATAATCTCAAACAGCTGAACCGCATGAGCGTACCGGATCAGGAGATGATGATCGGCCGTACCAAAGAAGCCAACGAAGAGATCGATGGCGACGAGCGTCCGGTCACGTCGCACCTGAGCCGCGTGGACTTAAAAGAAGATGGCAAAGGGCTGAAAATCGTCCGTCAGAGCCTGCCGTACGGCACCGCCAGCGGCACCCATGGTCTCTATTTCTGCGCCTACTGCGCGCGCCTGTATAACATCGAGCAGCAGCTGCTGAGCATGTTCGGCGATACCGACGGCAAACGCGACGCGATGCTGCGCTTCACTAAACCGGTGACCGGTGGCTATTACTTCGCGCCATCCCTGGAGCGTATTCAGGCGCTGTAA |
| NADH peroxidase | ATGAAAATCATTATCATAGGCGGGTCCTTTGGGGGTGTCAGCTGCGCCCGTGAAGCACGCCGATTATACCCTGACGCAGAGATTTTATTGATAGAAAAAAAAGTCCATCTTGGGTTTGTTCCAAGCGGTTTGTTATTGCTGATCGAAGGCAGGATCGCATCGCTGGAAGAAGCATTCTTTTGTACGAAGGAGCAGCTGGAGGCAGAAACGATCGACGTTGCCTTAGAAGAAACGCTTCTGGAGATTTTTCCTTCGGATCATCGGATTCGAACGGATAAGCGAGAACTAAGTTATGATCGACTCGTCTTGGCAGCAGGCTCCAGTCAAGATTCCACTGTTTTGCCACAACCAGAAGAAGCACTACTGACCTACAAGGAATATGAGGCAGCCAAGCATTTCTTGCAGCAATTGCCCCAAGCAGAGTCGATTACCATCGTAGGTGCTGGTCAAGCGGGGATGGAAGCGGCGAATACCTTGACTGCGATCGGAAAAAAAGTCAGTGTGATCGAATCAATGTCATATCCTTTATTCAAGTACTTTGATCACGATTTTTTGCAGCCCTTTTTAACGGAGATCGAAAAAATCCCCAACTTGCAGATGCATTGGTCCAAGCCAACTTTGGCAGTCGTTAAAACGGAAACAGGCTTTCGGATCGAAACCGCCGATCAAGGCTATGAAAGCGATCTGGTATTGACGACAGTCAACGTTCATCCTCGCTTGCCAGAGGCATTTGCGGTCTTTGAATTACACAGTGATCAAACCATCTGGACAGATGCCTACTTGGAAACTTCTGCCACGGATATTTTTGCGGTCGGCGATTTGATCCAAATTCCGAATGCTATTACGAAAGACACGGTCTATATGCCCTTAGTCAACAATGCTGTTCGCTCAGGAATCGCAGCAGCTCGTAATCTGGCAAGCAAAACCACCCCGTTTCGTGGCGGATTGCGGACAGTCGGAACGCAACTGTTTGGGTGGTACTTAGCAAGCACCGGCTTGACCGAGTCGGATCGGTTTTTCTTCGAATCTCCGATCGTCTGTCATGCGTTTACTGCCGCCAGTTCCTTGTTTGATCCAACACAGGTTCATGGAAAAGTGATGATCGAACAAGCCAGCGGCCGAATCGTTGGTGCGCAGCTTTTATCAAAAGCCAATATTTTGGAAAAAATCAATTTACTAGCGTTTGCGATCGAGCAGCAGGCAACGATGGAGGAGTTGACCCAAAAAGACTTTTTCTTCCATCCACGGTTCACTAATGTAATCGACGAAACTTTTCTATGGTCGGGAAGTGATGCCGATGAGACTTGA |

### Fig. S1 Phylum-level relative abundance of samples from four provinces

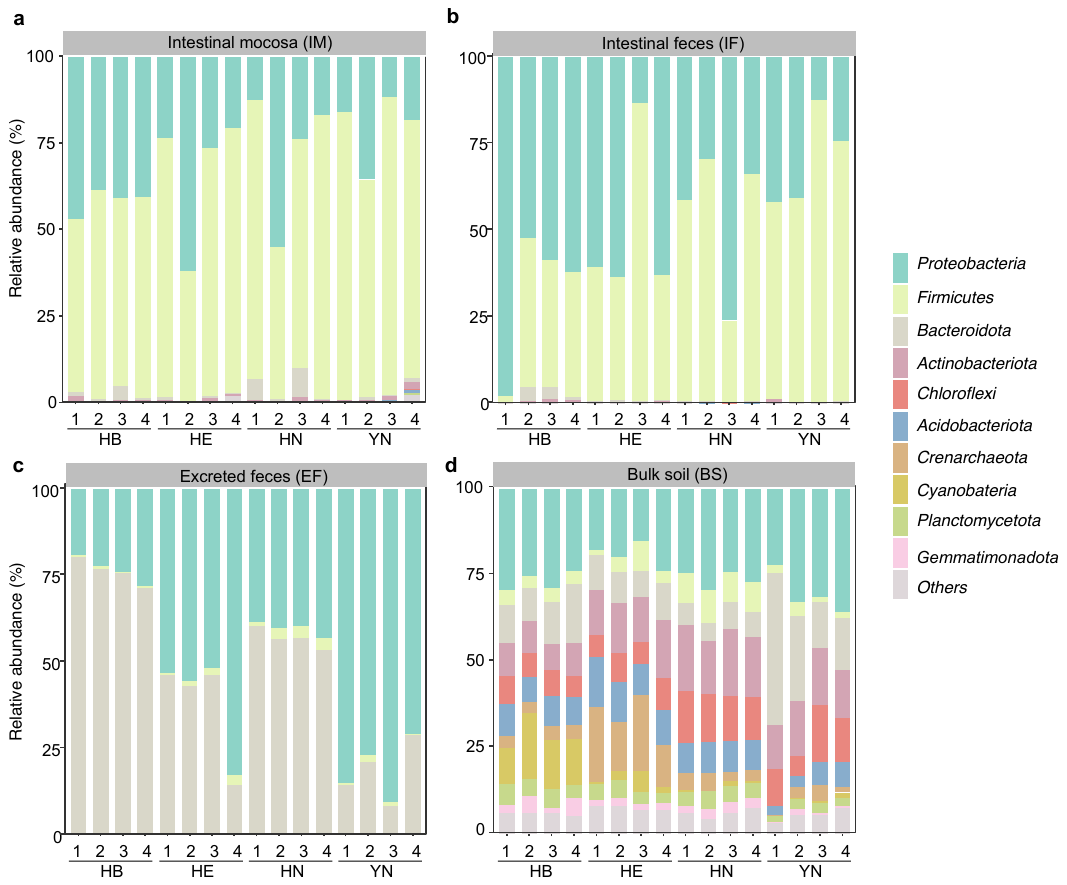

**Fig. S1** Phylum relative abundance of samples from four provinces collected in the **a**. Intestinal mucosa (IM), **b**. Intestinal feces (IF), **c**. Excreted feces (EF), **d**. Buk soil (BS).

**Fig. S2 Alpha-diversity of samples from four provinces**

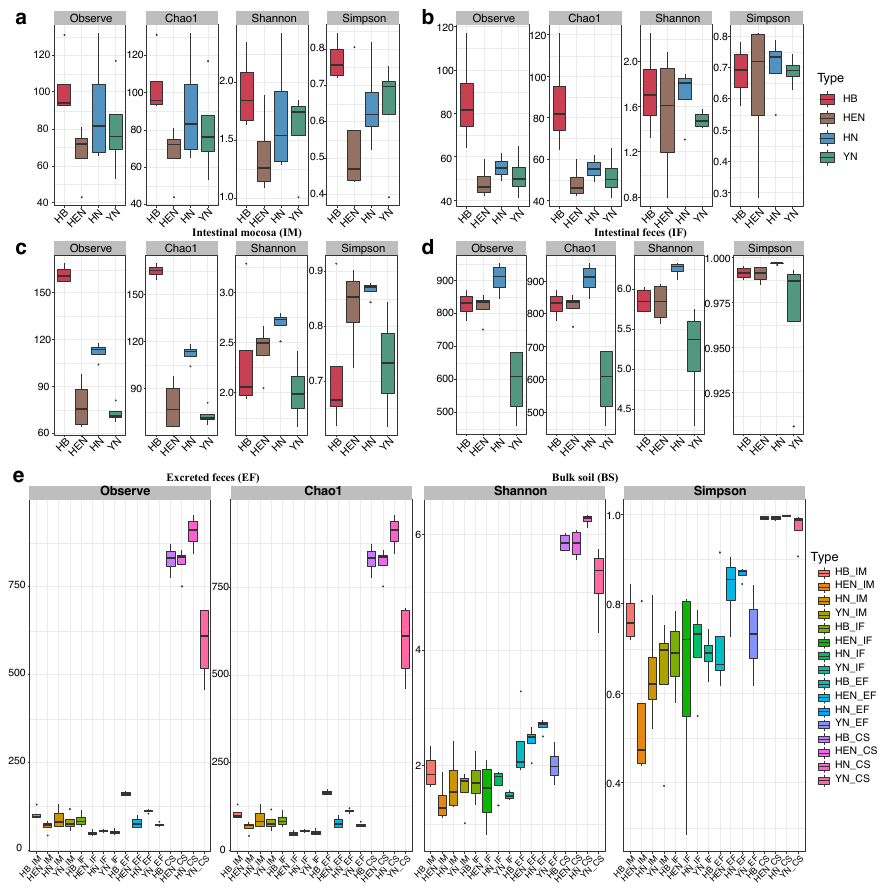

Alpha-diversity of samples from four provinces collected in **a**. Intestinal mucosa (IM), **b**. Intestinal feces (IF), **c**. Excreted feces (EF), **d**. Buk soil (BS), **e**. all sites including IM, IF, EF, and BS

**Fig. S3 Beta diversity of samples from four provinces collected in four sites.**

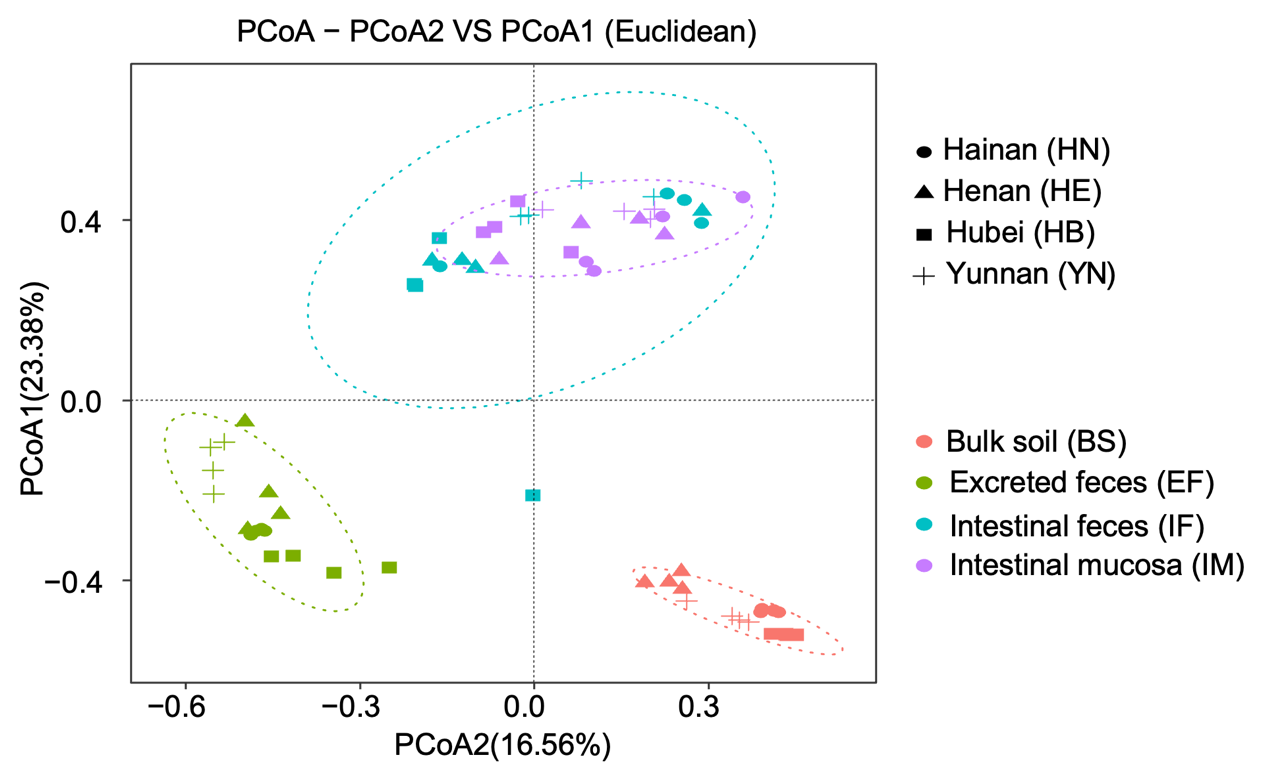
**Fig. S4 The Degradation characterization of PVC and composition of intestinal microbes after larvae ingested PVC.**

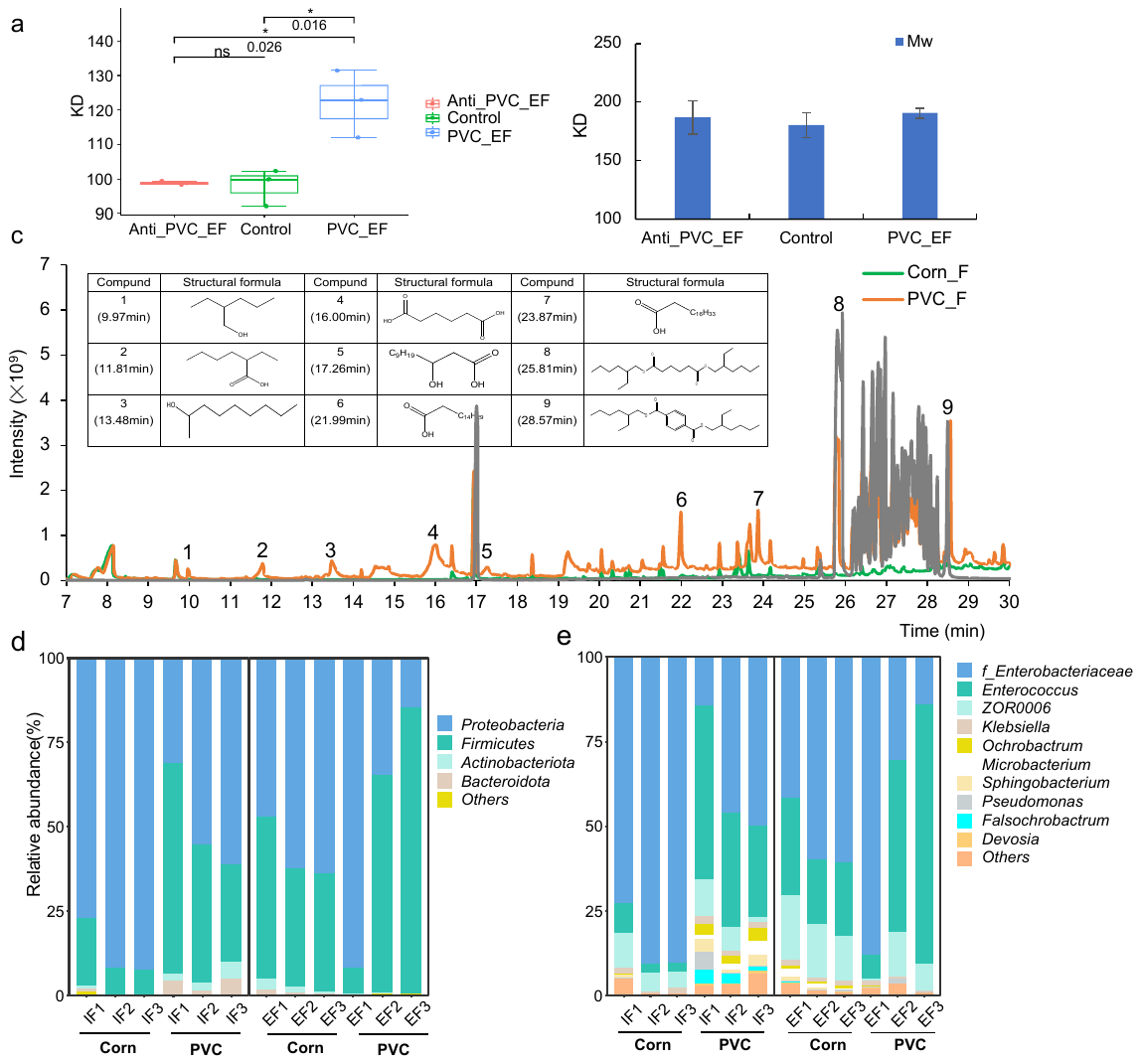

**a**, The detection results of the Mn of PVC film in different treatment groups*.* **b**, The detection results of the Mw of PVC film in different treatment groups. **c**, the possible degradation products of PVC film by the gut microbes. **d**, the composition of gut microbes after larvae ingested PVC.

### Fig. S5 Overlapped microbiota and differential microbiota between samples in S. frugiperda study

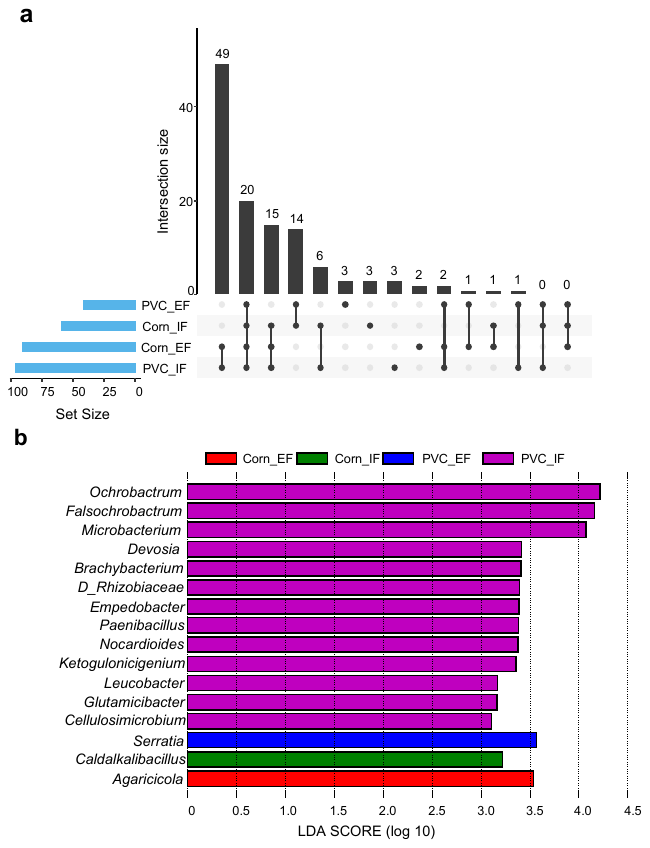

a. Overlapped microbiota between samples in *S. frugiperda*.

b. Differential microbiota in *S. frugiperda* study, Lefse was used to perform this statistic analysis (cutoff: LDA(log 10) > 3, p < 0.05).

### Fig. S6 Functional verification results of potential PVC-degrading strains and enzymes

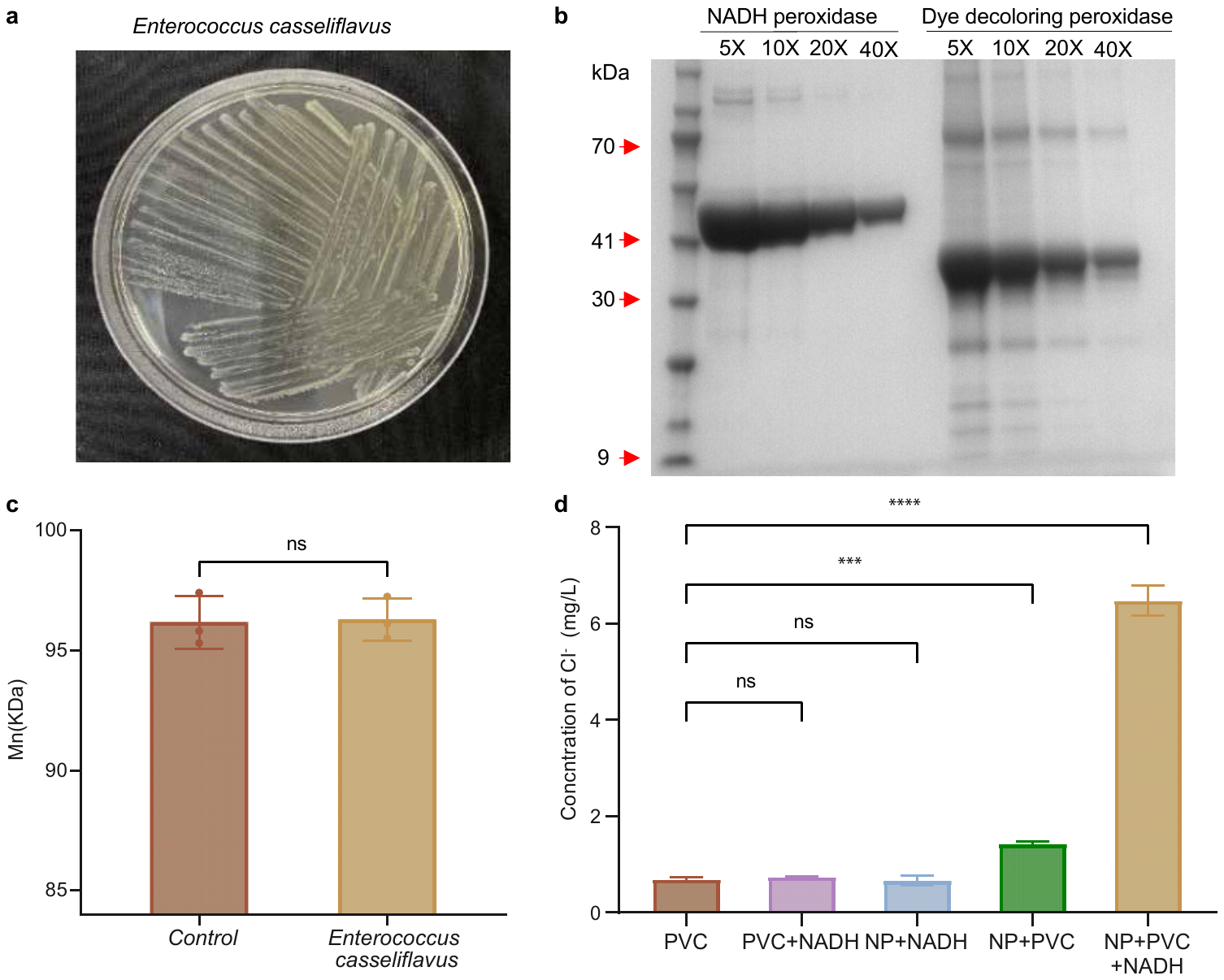

a, Growth morphology of PVC degrading *E. casseliflavus* EMBL-3 on LB solid medium.

b, Results of expression and purification of proteins of NADH peroxidase and dye decoloring peroxidase.

c, Molecular weight (Mn) of PVC film in the control group and the treatment group after 42 days (n = 3)

d, Dechlorination of control groups and enzyme treatment groups (n = 3; Significance, p < 0.01 indicated by **, p < 0.001 indicated by ***, p < 0.0001 indicated by ****).
